# Asymmetric chromatin capture and nuclear envelopes separate chromosomes in fused cells during mitosis

**DOI:** 10.1101/2020.12.17.423124

**Authors:** Bharath Sunchu, Nicole Lee, Roberto Carlos Segura, Chantal Roubinet, Clemens Cabernard

## Abstract

Hybrid cells derived through fertilization or somatic cell fusion recognize and separate chromosomes of different origin. The underlying mechanisms are unknown but could prevent aneuploidy and tumor formation. Here, we acutely induce fusion between *Drosophila* neural stem cells (neuroblasts; Nbs) and differentiating ganglion mother cells (GMCs) *in vivo* to define how epigenetically distinct chromatin is recognized and segregated. We find that Nb-GMC hybrid cells align both endogenous (neuroblast-origin) and ectopic (GMC-origin) chromosomes at the metaphase plate through centrosome derived dual-spindles. Mixing of endogenous and ectopic chromatin is prevented through an asymmetric, microtubule-dependent chromatin capture mechanism during interphase and physical boundaries imposed by nuclear envelopes. Although hybrid cells fail to accurately segregate ectopic chromatin, hybrid cells neither reduce the lifespan nor form visible tumors in host flies. We propose that Nb-GMC derived hybrid cells utilize asymmetric centrosome activity in interphase and nuclear envelopes to physically separate epigenetically distinct chromatin.

## Introduction

Dividing cells equally distribute the replicated chromosomes between the two sibling cells through microtubule-dependent attachment and segregation mechanisms (Maiato et al., 2017; McIntosh, 2016). Microtubules of bipolar spindles are connected to chromosomes via kinetochore proteins, which are localized on centromeric DNA (Thomas et al., 2017; Yu et al., 2019). Mitotic metazoan cells usually only form a single bipolar spindle, but certain insect species, arthropods or mouse zygotes form two distinct mitotic spindles (dual-spindles hereafter), which physically separate the maternal from the paternal chromatin in the first division after fertilization (Kawamura, 2001; Reichmann et al., 2018; Snook et al., 2011). Chromosome separation also occurs in hybrid cells, derived from somatic cell-cell fusion events (Heasley et al., 2017; Rieder et al., 1997). Dual-spindle dependent chromosome separation suggests the presence of specific chromosome recognition mechanisms, distinguishing between epigenetically distinct chromatin. The molecular nature of these recognition mechanisms is not known but could entail asymmetries in centromere binding proteins, kinetochore size or kinetochore composition (Akera et al., 2019; Arco et al., 2018; Drpic et al., 2018; Ranjan et al., 2019).

Over a century ago, unregulated cell-cell fusion between different somatic cells has been proposed to initiate tumor formation (Aichel, 1911). Aichel’s cell fusion model has the advantage that it can readily explain aneuploidy, a feature frequently observed at the early stages of tumor development (Ogle et al., 2005; Platt and Cascalho, 2019). Tetraploidy and supernumerary centrosomes – the natural products of cell fusion – predispose cells to aneuploidy through chromosome rearrangements (Fujiwara et al., 2005). Aichel’s cell fusion model still remains to be experimentally validated, which requires a detailed characterization of chromosome dynamics in fused cells *in vivo.*

Here, we ask how hybrid cells derived through cell-cell fusion of molecularly distinguishable cell types accurately recognize, separate and segregate epigenetically distinct chromosomes. To this end, we acutely fused *Drosophila* neural stem cells (neuroblasts (NBs), hereafter) with differentiating ganglion mother cells (GMCs) in the intact larval fly brain to create hybrid cells containing both neuroblast and GMC chromosomes. Unperturbed *Drosophila* neuroblasts divide asymmetrically, self-renewing the neural stem cell while forming a differentiating GMC. Neuroblasts are twice the size of GMCs, express the transcription factor Deadpan (Dpn^+^) and divide asymmetrically with a rapid cell cycle. The smaller GMCs can also be identified based on Prospero (Pros^+^) expression and divide only once with a long cell cycle (Gallaud et al., 2017).

Neuroblasts are intrinsically polarized, consisting of an apically localized Par complex, which is connected to the Pins complex, composed of Partner of Inscuteable (Pins; LGN/AGS3 in vertebrates), Gai and Mushroom body defects (Mud; NuMA in vertebrates, Lin-5 in C. elegans) (Gallaud et al., 2017; Loyer and Januschke, 2020; Sunchu and Cabernard, 2020). The Pins complex regulates spindle orientation during mitosis and biased centrosome asymmetry in interphase, manifested in the establishment and maintenance of an apical interphase microtubule organizing center (MTOC). The active interphase MTOC retains the daughter-centriole containing centrosome close to the apical cell cortex and pre-establishes spindle orientation in the subsequent mitosis (Gallaud et al., 2020; Gambarotto et al., 2019; Januschke et al., 2013, 2011; Januschke and Gonzalez, 2010; Rebollo et al., 2007; Rusan and Peifer, 2007).

We found that hybrid cells derived from such Nb-GMC fusions independently align the endogenous (neuroblast-origin) and ectopic (GMC-origin) chromosomes at the metaphase plate. We propose that these hybrid cells utilize asymmetric centrosome activity in interphase to capture, and nuclear envelopes to physically separate, epigenetically distinct chromatin *in vivo.* These findings provide mechanistic insight into how metazoan cells recognize and isolate chromosomes of different origin.

## Results

### Nb-GMC hybrid cells independently align Nb and GMC chromatin at the metaphase plate

To quantitatively describe chromosome dynamics in hybrid cells we developed an acute cell-cell fusion method in intact larval fly brains. We used a 532nm pulsed laser to induce a small lesion at the Nb – GMC interface, causing the GMC chromatin to enter the neuroblast cytoplasm. Neuroblasts can be distinguished from GMCs based on their size, molecular markers and cell cycle length (Figure 1 – figure supplement 1A). Targeted mitotic neuroblasts often retained the GMC chromosomes, creating a large apical hybrid cell containing one Dpn^+^ and one Pros^+^ nucleus (Nb – GMC hybrid). Most Nb – GMC hybrid cells normally localized the contractile ring marker non-muscle Myosin to the cleavage furrow and completed cytokinesis (Figure 1 – figure supplement 1B-E). Acute cell fusion can also result in the expulsion of the GMC chromatin, forming GMC – GMC hybrids (see Supplementary Figure 3c in (Roubinet et al., 2017)). Here, we are focusing on Nb – GMC hybrids (hybrid cells, hereafter) only.

To better characterize the dynamics of neuroblast (endogenous) and GMC (ectopic) chromosomes during mitosis, we induced cell fusion at different cell cycle stages in wild type neuroblasts, expressing the canonical chromosome marker His2A::GFP. We hypothesized that hybrid cells derived from Nb-GMC fusions early in the cell cycle could (1) align only the neuroblast chromosomes at the metaphase plate, (2) congress a mix of neuroblast and GMC chromosomes or (3) separate and align the two chromosome pools at the metaphase plate (Figure 1A). We found that the endogenous and ectopic chromatin was separated and distinguishable when fusions were induced in early mitosis. Both the ectopic and endogenous chromatin aligned at the metaphase plate (Figure 1B; video 1&2). Nb-GMC fusions could be induced at all cell cycle stages but GMC chromosomes aligned at the metaphase plate more accurately in hybrid cells derived from interphase or early prophase fusions (Figure 1C, D). All fusions reported here were performed with non-mitotic GMCs. We conclude that hybrid cells derived from fusions between interphase Nbs and non-mitotic GMCs accurately align ectopic and endogenous chromatin at the metaphase plate.

**Figure 1:**
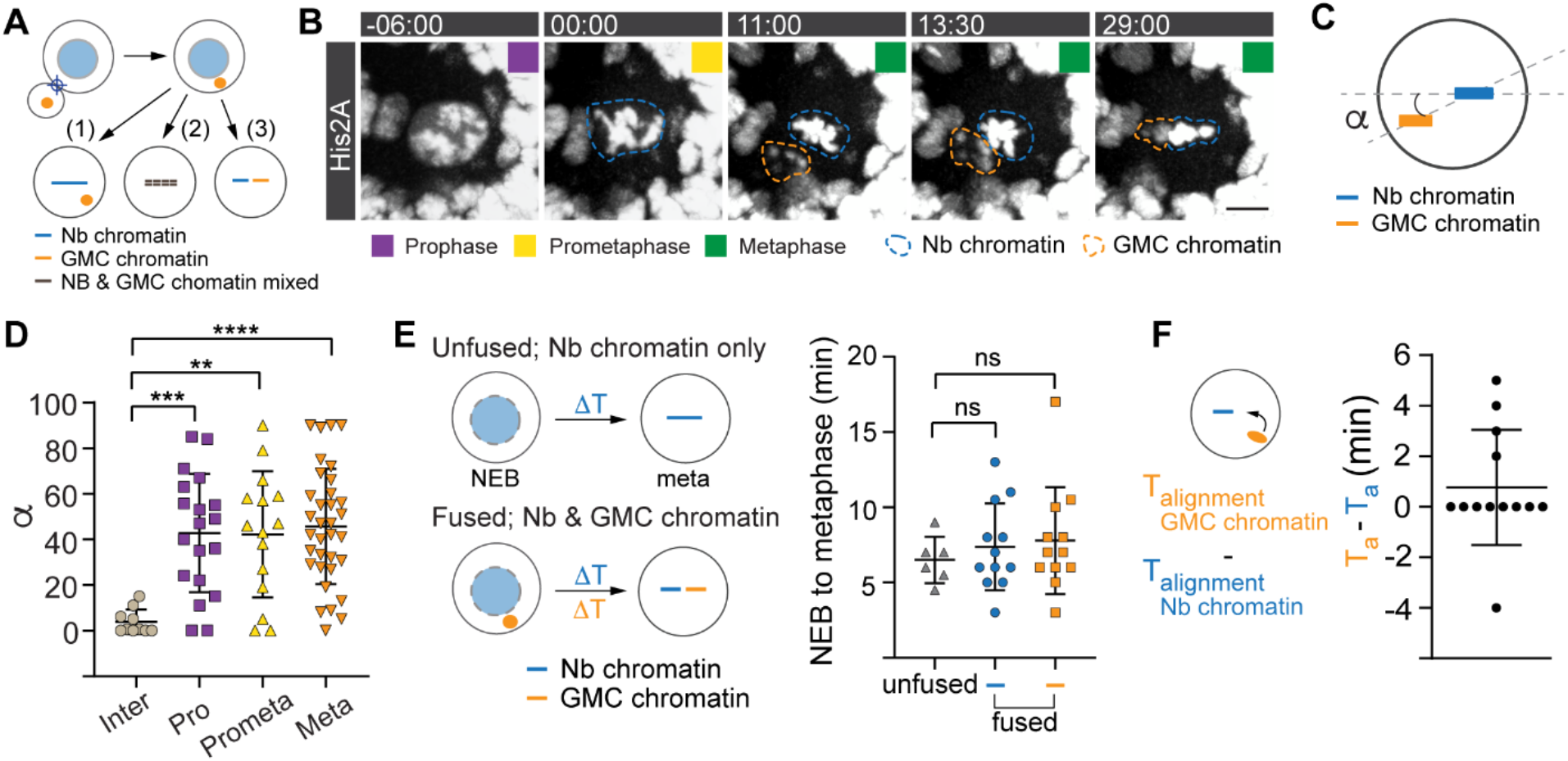
Nb-GMC hybrid cells align Nb and GMC chromatin independently at the metaphase plate. **(A)** Potential outcomes of Nb-GMC fusions: Nb – GMC derived hybrid cells could (1) only align neuroblast chromosomes, (2) congress a mix of endogenous and ectopic chromosomes or (3) separately align Nb and GMC chromosomes at the metaphase plate. **(B)** Representative image sequence of a dividing third instar larval Nb-GMC hybrid cell obtained from an interphase fusion, expressing the histone marker His2A::GFP (dashed blue circle; endogenous chromatin, dashed orange circle; ectopic chromatin). **(C)** Alignment of endogenous and ectopic chromatin in hybrid cells derived from interphase (inter), prophase (pro), prometaphase (prometa) or metaphase (meta) fusions were quantified with angle measurements in metaphase and plotted in **(D)**. **(E)** Chromosome alignment time for endogenous (blue circles) and ectopic chromosomes (orange squares) compared to unfused control neuroblasts (grey triangles). **(F)** Scatter plot showing the time difference (Ta; time of alignment) between endogenous and ectopic chromatin. Colored boxes represent corresponding cell cycle stages. One-way ANOVA was used for (D) and (E). Error bars correspond to standard deviation (SD). * p<0.05, **p<0.01, **** p<0.000.1. For this and subsequent figures, exact p values and complete statistical information can be found in the Extended data table 1. Time in mins:secs. Scale bar is 5 μm.

We next asked whether GMC chromatin congresses independently of neuroblast chromatin. To this end, we measured the time between nuclear envelope breakdown (NEB) and chromosome alignment at the metaphase plate for Nb and ectopic GMC chromatin in hybrid cells derived from interphase and early prophase fusions (Figure 1E). In most hybrid cells, ectopic and endogenous chromatin was distinguishable based on differences in location and intensity (see Figure 1B; video 1&2). In untargeted control neuroblasts (unfused), Nb chromosomes aligned at the metaphase plate within 6.5 minutes after NEB (SD = 1.55; n = 6), which is insignificantly faster than the neuroblast chromosomes of hybrid cells (t = 6.9 mins; SD = 2.84; n = 11). GMC chromosomes aligned within 7.63 mins (SD = 3.69; n = 11), statistically not significantly different from unperturbed wild type chromosomes (Figure 1E). In most Nb-GMC hybrids, the endogenous neuroblast and the ectopic GMC chromosomes aligned at the metaphase plate with no significant time difference. However, in a few cases, ectopic chromatin aligned before or after the neuroblast chromatin (Figure 1F). These results suggest that the neuroblast and GMC chromatin can move independently to the metaphase plate in hybrid cells.

### Ectopic spindles distinguish between Nb and GMC chromatin

We next investigated the mechanisms underlying independent Nb/GMC chromosome alignment, considering the following possibilities: ectopic chromosomes could be aligned together with the endogenous chromosomes via a single bipolar spindle. Alternatively, hybrid cells could form multiple bipolar spindles, which attach to either the neuroblast’s, GMC’s, or chromosomes from both cell types (Figure 2A). Live cell imaging showed that hybrid cells derived from interphase Nb-GMC fusions contained double spindles in almost all cases, whereas the vast majority of cell fusions induced in metaphase only formed one mitotic spindle (Figure 2B, C & video 3). Most hybrid cells contained two mitotic spindles, which typically formed at the same time (Figure 2D, E). We quantified spindle alignment and positioning dynamics by measuring the angle and distance between the two spindles during metaphase (Figure 2F and methods). The two spindles were initially misaligned and separated but decreased their inter-spindle distance and angle during metaphase (Figure 2G-I). We conclude that Nb-GMC hybrid cells align neuroblast and GMC chromosomes separately at the metaphase plate through the formation of independent mitotic spindles.

**Figure 2:**
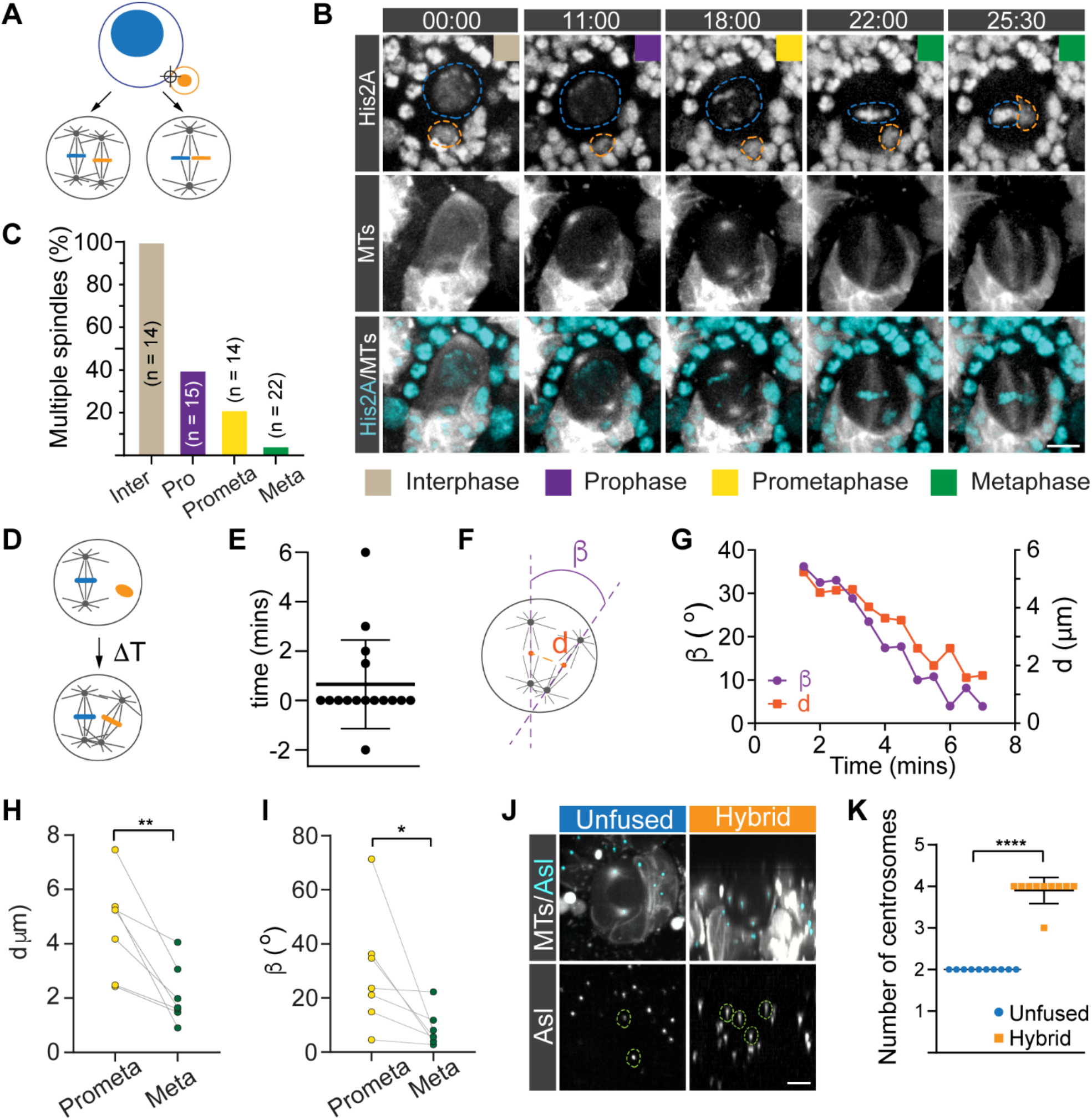
Dual spindles align endogenous and ectopic spindles separately at the metaphase plate. **(A)** Hypothetical outcomes of spindle organization after Nb-GMC fusions: hybrid cells could align neuroblast and GMC chromosomes either through a single or dual-spindle mechanism. **(B)** Representative third instar larval Nb-GMC hybrid cell, expressing the histone marker His2A::GFP (white in top row; cyan in merged channel below) and the MT marker cherry::Jupiter (white in middle and bottom row). 00:00 refers to nuclear envelope breakdown (NEB). **(C)** Quantification of hybrid cells containing dual- or multiple spindles for hybrid cells derived from interphase (inter), prophase (pro), prometaphase (prometa) or metaphase (meta) fusions. **(D)** The time difference between endogenous and ectopic spindle formation was measured in hybrid cells and plotted in **(E)**. **(F)** Spindle angle and inter-spindle distances were measured during mitosis. A representative example is shown in **(G)**. Quantification of inter-spindle **(H)** distances and **(I)** angles at prometaphase and metaphase. **(J)** A representative third instar larval Nb-GMC hybrid, expressing the centriole marker Asl::GFP (cyan; top row. White; bottom row) and the spindle marker cherry::Jupiter (MTs; white in top row). Centrosomes were highlighted with green dashed circles. **(K)** Comparison of centrosome number between unfused wild type and hybrid cells. Colored boxes represent corresponding cell cycle stages. Error bars correspond to SDs. Figures (H) and (I); two-sided paired t-test, figures (K); two-sided unpaired t-test. * p<0.05, **p<0.01, **** p<0.000. Time in mins:secs. Scale bar is 5 μm.

### Ectopic spindles are nucleated from GMC centrosomes

Mitotic spindles can be nucleated through the centrosome-dependent, chromatin or microtubule pathway but when centrosomes are present, the centrosome pathway prevails (Prosser and Pelletier, 2017). To elucidate the mechanisms underlying ectopic spindle formation, we induced Nb-GMC fusions in interphase wild type neuroblasts expressing live centriole (Asterless; Asl::GFP) and spindle (cherry::Jupiter) markers, and assayed centrosome dynamics and spindle formation throughout mitosis. Normal wild type neuroblasts usually contained two Asl::GFP positive centrioles in mitosis, forming a single bipolar spindle. However, in Nb-GMC hybrids, we predominantly found four Asl::GFP positive centrioles, two of which were introduced from the GMC (Figure 2J,K). The GMC centrosomes nucleated an ectopic bipolar spindle that subsequently aligned with the main neuroblast spindle. Although multipolar spindle formation can be prevented through centrosome clustering (Quintyne et al., 2005), we often observed that GMC-derived and Nb-derived spindles were oriented in parallel to each other but remained separate. These data suggest that ectopic spindles are formed through the centrosome pathway, using centrioles originating from GMCs.

### Microtubule-dependent, asymmetric chromatin-centrosome attachments retain chromosomes close to the apical neuroblast cortex during interphase

Our data suggest that Nb and GMC chromatin are being separated through an endogenous, Nb-derived and an ectopic, GMC-derived mitotic spindle. We next investigated how these spindles distinguish between Nb and GMC chromosomes. During mitosis, microtubules emanate from centrosomes and attach to sister chromatids via kinetochore proteins, localizing to the centromeric region (Fukagawa and Earnshaw, 2014). *Drosophila* male germline stem cells contain asymmetric levels of the centromere-specific H3 variant (Centromere identifier (Cid) in flies (Henikoff et al., 2000) (Ranjan et al., 2019), prompting us to investigate whether hybrid cell spindles differentiate between endogenous and ectopic chromosomes based on differing levels of Cid. We induced Nb-GMC fusions of wild type cells expressing Cid::EGFP (Ranjan et al., 2019) in interphase or early prophase and measured Cid intensity on both GMC and Nb chromatin. These measurements did not reveal a significant intensity difference between Nb and GMC Cid (Figure 3 – figure supplement 1A). However, we noticed that endogenous Cid::EGFP was localized in very close proximity to the apical centrosome in unperturbed interphase and prophase wild type neuroblasts (Figure 3A & video 4). Cid::EGFP remained associated with chromatin throughout the neuroblast cell cycle, excluding the possibility that early Cid clusters are not connected with chromatin (Figure 3 – figure supplement 1B & video 5).

**Figure 3:**
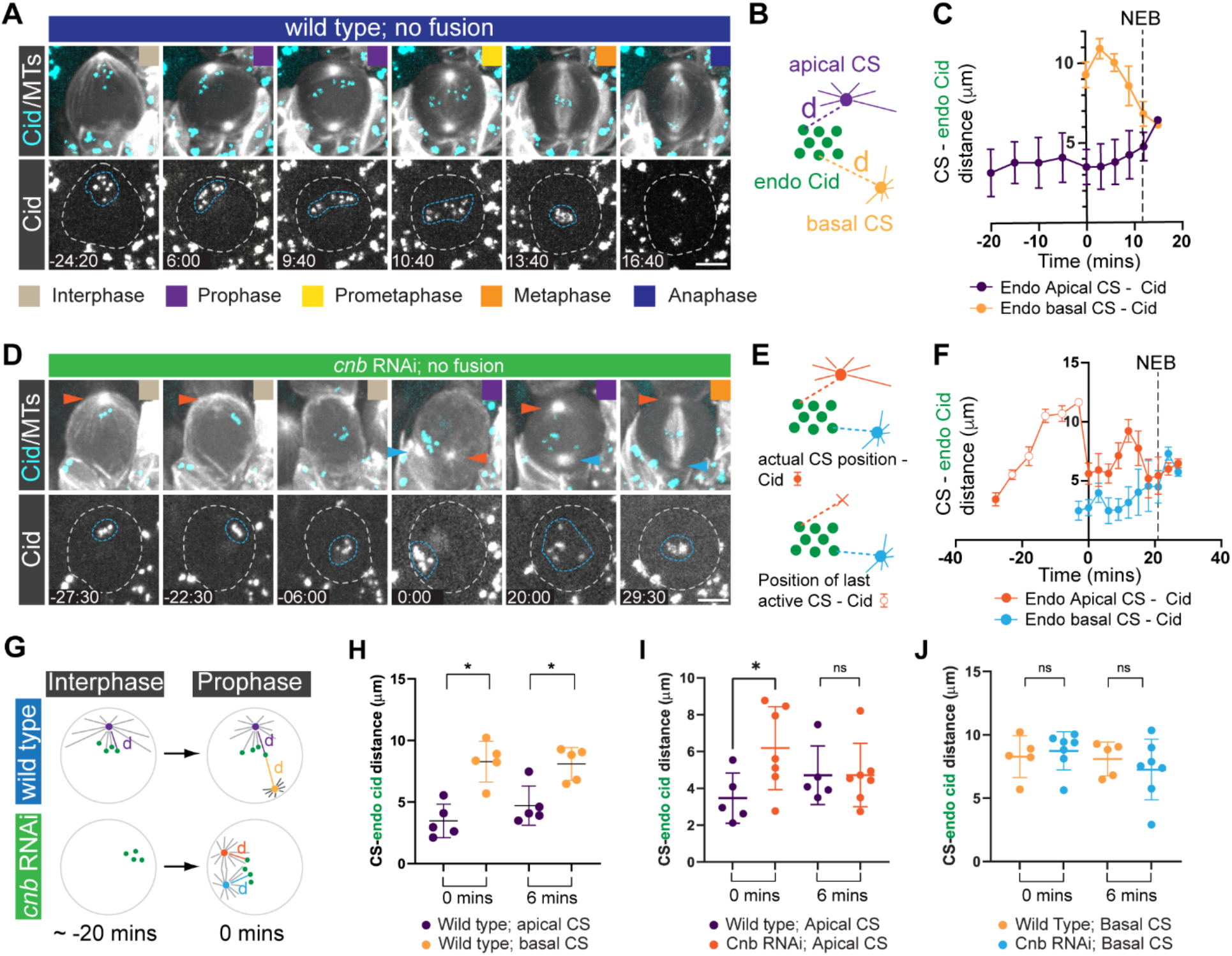
Biased MTOC activity retains Cid in the apical neuroblast hemisphere during interphase and early mitosis. **(A)** Representative third instar larval neuroblast expressing the centromere specific Histone-3 variant marker, Cid::EGFP (cyan; top, white; bottom) and the microtubule marker cherry::Jupiter (white; top). Colored boxes represent corresponding cell cycle stages. **(B)** The distance (purple and yellow dashed lines) between the apical (purple) and basal (yellow) centrosome, and individual Cid clusters (green circles) were measured throughout the cell cycle and plotted in **(C)**. **(D)** Representative third instar larval neuroblast expressing *cnb* RNAi, cherry::Jupiter (white; top) and Cid::EGFP (cyan; top, white; bottom row). Red arrowheads highlight the apical MTOC. Blue arrowheads highlight the maturing basal MTOC. The blue dashed circle highlights Cid clusters. The cell outline is indicated with the white dashed. ‘Apical’ centrosome refers to the centrosome destined to be positioned on the apical cortex, whereas ‘basal’ centrosome will be inherited by the basal GMC. **(E)** CS – Cid distance measurements were performed in *cnb* RNAi expressing Nbs. Once the apical CS disappeared in interphase, the last detectable position was used as a reference point (orange cross; open circles). **(F)** CS – Cid measurements for *cnb* RNAi expressing Nbs. Closed arrows refer to actual CS – Cid measurements. Open circles denote Cid – previous active CS measurements. **(G)** Wild type neuroblasts maintain apical CS – Cid attachments in prophase, due to asymmetric MTOC activity and microtubule-dependent interphase centrosome – Cid attachments. *cnb* RNAi expressing neuroblasts lose MTOC activity in interphase, randomizing the position of Cid clusters. When centrosomes mature again in prophase, both centrosomes simultaneously attach to Cid clusters. **(H)** Centrosome – Cid distance of an unperturbed wild type neuroblast at the time of basal centrosome maturation (0 mins) and 6 mins thereafter. **(I, J)** Cid – centrosome distance measurement in *cnb* RNAi expressing neuroblasts. Error bars correspond to SDs. Figure (H, I, J); two-sided paired t-test. ns; no signficiance. * p<0.05. Time in mins:secs. Scale bar is 5 μm.

Interphase wild type neuroblasts contain only one active apical microtubule organizing center (MTOC), which retains the daughter centriole-containing centrosome close to the apical neuroblast cortex. The mother-centriole-containing centrosome is inactive in interphase but matures from prophase onward, positioning itself on the basal cell cortex (Gallaud et al., 2020; Januschke et al., 2013, 2011; Januschke and Gonzalez, 2010). We measured the distance of individual Cid::EGFP clusters to the apical and basal centrosome in unperturbed wild type neuroblasts and found that during interphase Cid was always in close proximity to the apical centrosome. After nuclear envelope breakdown (NEB), Cid moved progressively towards the metaphase plate. Once the basal centrosome appeared (0 mins), Cid was still closer to the apical than the basal centrosome and this distance asymmetry was also observed 6 minutes after the appearance of the basal centrosome (Figure 3B, C, H).

The proximity of Cid::EGFP clusters to the active interphase MTOC suggests a microtubule-dependent chromosome attachment mechanism. Indeed, wild type neuroblasts treated with the microtubule-depolymerizing drug colcemid showed a strong correlation between apical MTOC activity and Cid localization; as MTs depolymerized after colcemid addition, Cid progressively moved away from the apical cortex towards the cell center (Figure 3 – figure supplement 1C, D & video 6). To test whether MTOCs are connected to Cid clusters during interphase, we removed the centriolar protein Centrobin (Cnb; CNTROB in humans). Neuroblasts lacking Cnb fail to maintain an active apical interphase MTOC but regain normal MTOC activity during mitosis (Januschke et al., 2013). Neuroblasts expressing *cnb* RNAi lost apical Cid localization after the apical centrosome downregulated its MTOC activity. However, maturing centrosomes reconnected with Cid in prophase (Figure 3D-G & video 7). Cid’s proximity to the apical MTOC (‘apical’ refers to the centrosome destined to move to the apical cortex) in *cnb* RNAi expressing neuroblasts was much more varied compared to wild type. At 6 minutes after centrosome maturation onset, Cid – apical MTOC distance was comparable to wild type, as were Cid – basal centrosome distance relationships (Figure 3I, J). This suggests that when centrosomes mature during mitosis and MTOC activity is restored, Cid – MTOC attachments can be reestablished during early mitosis in *cnb* RNAi expressing neuroblasts (Figure 3G). We conclude that in wild type Nbs, Cid-containing chromatin is already attached to the apical centrosome prior to entry into mitosis.

### Asymmetric centrosome-chromatin attachments contribute to the separation of endogenous and ectopic chromosomes

Based on these observations, we hypothesized that the separation between endogenous and ectopic chromosomes could be due to a pre-attachment mechanism, preventing mixing of Nb and GMC chromosomes. To test this hypothesis, we first analyzed Cid localization in relation to the endogenous and ectopic centrosomes in wild type hybrid cells. Similar to unperturbed wild type neuroblasts, endogenous Cid is also localized in close proximity to the endogenous apical centrosome in wild type hybrid cells (Figure 4A-C, G, H & video 8; ‘apical’ refers to the centrosome destined to segregate into the large apical sibling cell). Ectopic Cid, however, appeared closer to ectopic centrosomes (Figure 4A-C, I & video 8; 0 mins refers to the appearance of the endogenous basal centrosome; ‘0’ refers to the appearance of the ectopic centrosomes).

**Figure 4:**
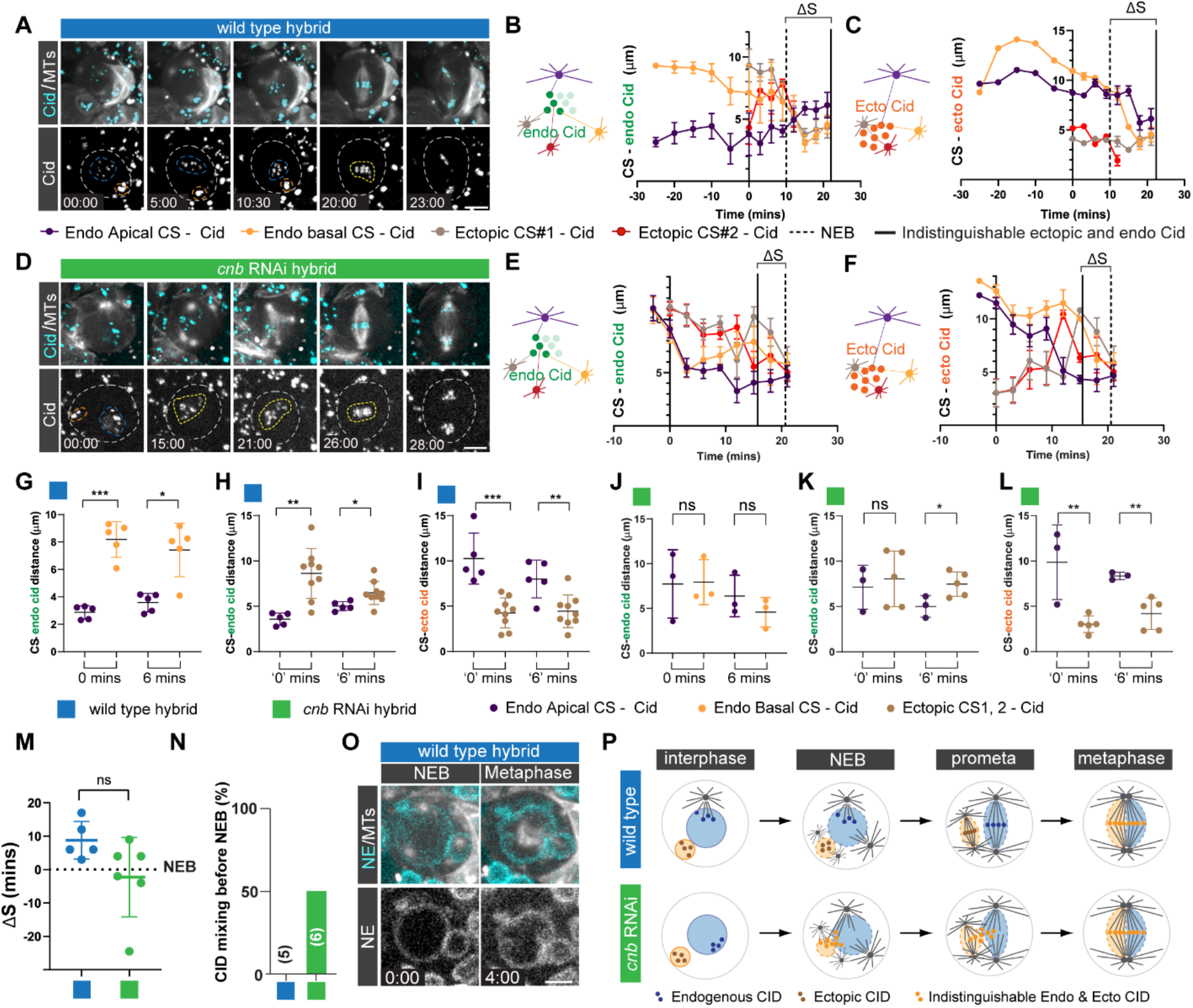
Asymmetric microtubule dependent centrosome-chromatin attachments contribute to the separation of endogenous and ectopic chromosomes. Representative third instar larval **(A)** wild type or **(D)** *cnb* RNAi expressing Nb-GMC hybrid cell, expressing Cid::EGFP (top row; cyan, bottom row; white) and the microtubule marker cherry::Jupiter (white; top row). Neuroblast-derived and GMC-derived Cid clusters are outlined with a blue and orange dashed line, respectively. The white dashed line labels the cell outline. Indistinguishable GMC and Nb Cid clusters are highlighted with yellow dashed circles. The distance of **(B)** endogenous or **(C)** ectopic Cid from the wild type hybrid cell shown in (A) in relation to GMC and Nb centrosomes (CS) plotted over time. **(E)** Endogenous or **(F)** ectopic Cid – centrosome distance measurements of the *cnb* RNAi expressing hybrid cell shown in (D). The vertical solid line indicates the time point when endogenous and ectopic Cid can no longer be distinguished. Vertical dashed line indicates nuclear envelope breakdown. For (B), (C), (E), (F): ?S refers to the time difference between NEB and when ectopic and endogenous CID becomes indistinguishable. Measurements of endogenous Cid – apical (purple) or basal (yellow) endogenous centrosomes were plotted when **(G)** the endogenous basal centrosome matured (0 mins) and 6 minutes thereafter (6mins) or **(H)** the ectopic centrosomes matured (‘0’ mins) and 6 minutes thereafter. Corresponding measurements of *cnb* RNAi expressing hybrid cells are shown in **(J)** and **(K)**. Distance of ectopic Cid in relation to endogenous and ectopic centrosomes shown for **(I)** wild type or (L) *cnb* RNAi expressing hybrid cells. **(M)** Time interval between NEB and mixing of endogenous and ectopic Cid for wild type (blue circles) and *cnb* RNAi expressing (green circles) hybrid cells. **(N)** Bar graph showing percentage of hybrid cells in which the endogenous and ectopic CID mixed prior to NEB. **(O)** A third instar wild type hybrid cell expressing the nuclear envelope (NE) marker, koi::GFP (cyan; top row, white; bottom row) and cherry::Jupiter. **(P)** Schematic summary: separation of endogenous and ectopic chromosomes involves microtubuledependent asymmetric chromatin-centrosome attachments and physical separation through nuclear envelopes. *cnb* RNAi expressing neuroblasts release endogenous Cid during interphase, partially randomizing re-attachment in the subsequent mitosis, which causes premature mixing of endogenous and ectopic Cid. Colored boxes refer to the indicated cell cycle stages. Error bars correspond to SDs. Figure (G); two-sided paired t-test and Figures (H-L); two-sided unpaired t-test. ns; no signficiance. * p<0.05, **p<0.01, ***p<0.0001, **** p<0.0001. Time in mins:secs. Scale bar is 5 μm.

We next attempted to randomize Cid – centrosome distance relationships by inducing fusions in *cnb* RNAi expressing neuroblasts, since loss of interphase MTOC activity released endogenous Cid from the apical centrosome (Figure 3D-J and Figure 3 – figure supplement 1C, D). In contrast to wild type hybrid cells, endogenous Cid is roughly equidistant to the endogenous and ectopic centrosomes in *cnb* RNAi expressing hybrid cells at 0 mins and ‘0’ mins respectively (Figure 4D-F, J, K & video 9). However, ectopic Cid was still closer to the ectopic centrosomes than to the endogenous apical centrosome in *cnb* RNAi expressing hybrid cells (Figure 4D, F, L & video 9). In both wild type and *cnb* RNAi expressing hybrid cells, ectopic and basal centrosomes were about equidistant to endogenous Cid, but ectopic centrosomes were closer to ectopic Cid (Figure 4 – figure supplement 1A-D). Taken together, we conclude that in hybrid cells the apical MTOC forms an asymmetric attachment to endogenous Cid-containing chromosomes prior to entry into mitosis.

### Nuclear envelopes separate ectopic and endogenous chromatin in hybrid cells

We next asked whether early MTOC – Cid attachments are sufficient to prevent endogenous and ectopic chromosome separation and tracked endogenous and ectopic Cid::EGFP after induced cell fusion. In wild type hybrid cells, endogenous and ectopic CID clusters started to congress at the metaphase plate and became difficult to clearly separate 8.8 mins (SD = 5.63; n = 5; Figure 4M) after NEB. However, 50% of *cnb* RNAi hybrid cells showed endogenous and ectopic Cid mixing prior to NEB (Figure 4D, N), although the time difference was not significantly different to wild type hybrid cells (Figure 4M).

Neuroblasts undergo semi-closed mitosis, mostly retaining a matrix composed of nuclear envelope proteins around the mitotic spindle (Katsani et al., 2008). We imaged hybrid cells with the nuclear envelope marker Klaroid, using the protein-trap line koi::EGFP (Buszczak et al., 2007). We confirmed that unperturbed wild type neuroblasts contain a nuclear envelope matrix surrounding the mitotic spindle during mitosis (Figure 4 – figure supplement 1E). Similarly, wild type hybrid cells contain two nuclear envelopes during mitosis (Figure 4O and Figure 4 – figure supplement 1F). Taken together, these data suggest that both asymmetric MTOC-Cid attachments in interphase and nuclear envelopes establish and maintain the physical separation between endogenous and ectopic chromosomes in hybrid cells. Loss of biased interphase MTOC activity removes asymmetric MTOC-Cid attachments and allows for cross-connections between endogenous centrosomes and ectopic Cid in early mitosis (Figure 4P).

### Hybrid cells segregate endogenous and ectopic chromosomes independently

We next investigated whether both bipolar spindles are functional in Nb-GMC hybrid cells. Erroneous or incomplete microtubule-kinetochore attachments trigger the spindle assembly checkpoint (SAC), preventing or delaying anaphase entry (Musacchio, 2015). Since the kinetochore-derived ‘wait anaphase’ signal is diffusible (Heasley et al., 2017), ectopic spindles should thus also obey the SAC in Nb-GMC hybrid cells. We tested whether hybrid cells contain functional microtubule-kinetochore attachments by measuring the time between finished chromosome alignment at the metaphase plate and chromosome separation in Nb-GMC hybrids expressing His2A::GFP and cherry::Jupiter (Figure 5A, B). Unperturbed control neuroblasts usually initiate anaphase onset within 2.63 minutes (SD=1.84; n=12) after chromosomes are aligned at the metaphase plate. In hybrid cells derived from interphase fusions, endogenous and ectopic chromatin entered anaphase 4.14 mins (SD= 2.23; n=14) and 5.12 minutes (SD= 1.85; n=13) after metaphase alignment. Only ectopic chromatin for prophase-induced hybrid cells showed a significantly delayed anaphase onset (Average: 10.25 mins; SD= 4.12; n=6) (Figure 5C). Ectopic chromatin never separated before endogenous chromosomes but entered anaphase with a few minutes’ delay (Figure 5D). We conclude that in Nb-GMC hybrids, endogenous and ectopic chromosomes establish correct MT-kinetochore attachments, thereby fulfilling the spindle assembly checkpoint necessary to enter anaphase. However, given the delays in ectopic chromosome separation, we further conclude that ectopic spindles can initiate chromatid separation independently from the endogenous neuroblast spindle.

**Figure 5:**
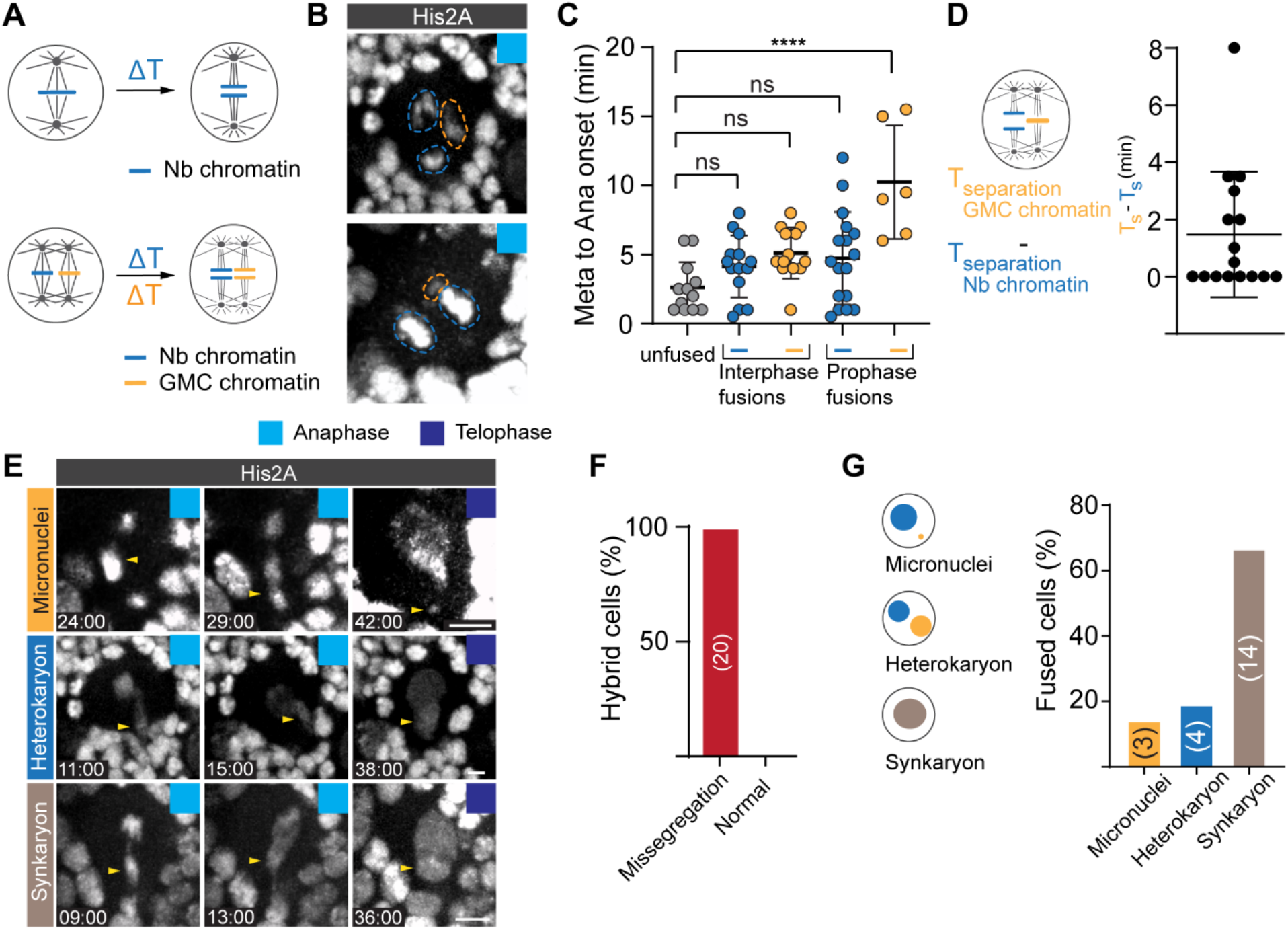
Ectopic chromosomes in hybrid cells segregate independently but erroneously. **(A)** The time between chromosome alignment (metaphase) and separation (anaphase onset) was measured for endogenous and ectopic chromosomes in wild type hybrid cells and plotted in (C). **(B)** Representative examples of delayed (top row) and simultaneous (bottom row) segregation of endogenous (blue dashed circle) and ectopic (orange dashed circle) chromosomes in wild type hybrid cells expressing the chromatin marker His2A::GFP. **(C)** Quantification of metaphase to anaphase onset in control Nbs, compared to interphase and prophase Nb-GMC hybrids. **(D)** Average time difference between Nb and GMC chromatin anaphase onset in Nb-GMC hybrid cells. **(E)** Representative third instar larval Nb-GMC hybrids expressing histone marker His2A::GFP showing missegregating chromatids (yellow arrowheads) during anaphase, resulting in micronuclei (top row), heterokaryon (middle row) or synkaryon (bottom row) formation. Time stamps are in relation to NEB (=0:00). **(F)** Bar graph showing percentage of Nb-GMC hybrid cells with missegregating chromatids. **(G)** Bar graph quantifying the percentage of fused cells with micronuclei, heterokaryons or synkaryons. ΔT; time difference. Error bars correspond to SDs. One way ANOVA test was used in (C). ns; no significance. **** p<0.0001. Time in mins:secs. Scale bar is 5 μm.

### Hybrid cells are insufficient to induce tumors in wild type host flies

Finally, we assessed the accuracy of chromosome segregation in wild type hybrid cells. Using the canonical chromosome marker His2A::GFP we detected chromosome missegregation – ranging from lagging chromosomes to chromosome bridges – in all wild type hybrid cells (Figure 5E,F & video 10). Chromosome segregation defects can result in aneuploidy and micronuclei formation (Molina et al., 2020). We found a small percentage of hybrid cells containing micronuclei, but more frequently discovered heterokaryons (hybrid cells containing two nuclei of different origins). In most cases, hybrid cells fused both nuclei into one, forming synkaryons (Figure 5G).

Aneuploidy has been proposed to be a hallmark of cancer (Molina et al., 2020) but appears to be context dependent (Ben-David and Amon, 2020). To test whether neuroblast – GMC derived hybrid cells are sufficient to induce tumor formation, we grafted His2A::GFP expressing larval fly brains after successful induction of cell fusion into the abdomen of wild type adult hosts (Rossi and Gonzalez, 2015) and monitored the host flies for tumor formation and life span changes. As previously reported (Caussinus and Gonzalez, 2005), brat RNAi expressing brains formed visible tumors in host flies by day 30 and caused a reduction in lifespan of the host. However, larval brains without attempted fusions (wild type transplants), with attempted but unsuccessful fusions (controlling for the effect of laser ablation), or with successful fusions, showed neither tumor growth by day 30 (or after), nor a reduction in lifespan (Figure 6A, B, C). We conclude that chromosome segregation is defective in Nb-GMC hybrid cells but insufficient to form visible tumors in otherwise normal and unperturbed larval fly brains.

**Figure 6:**
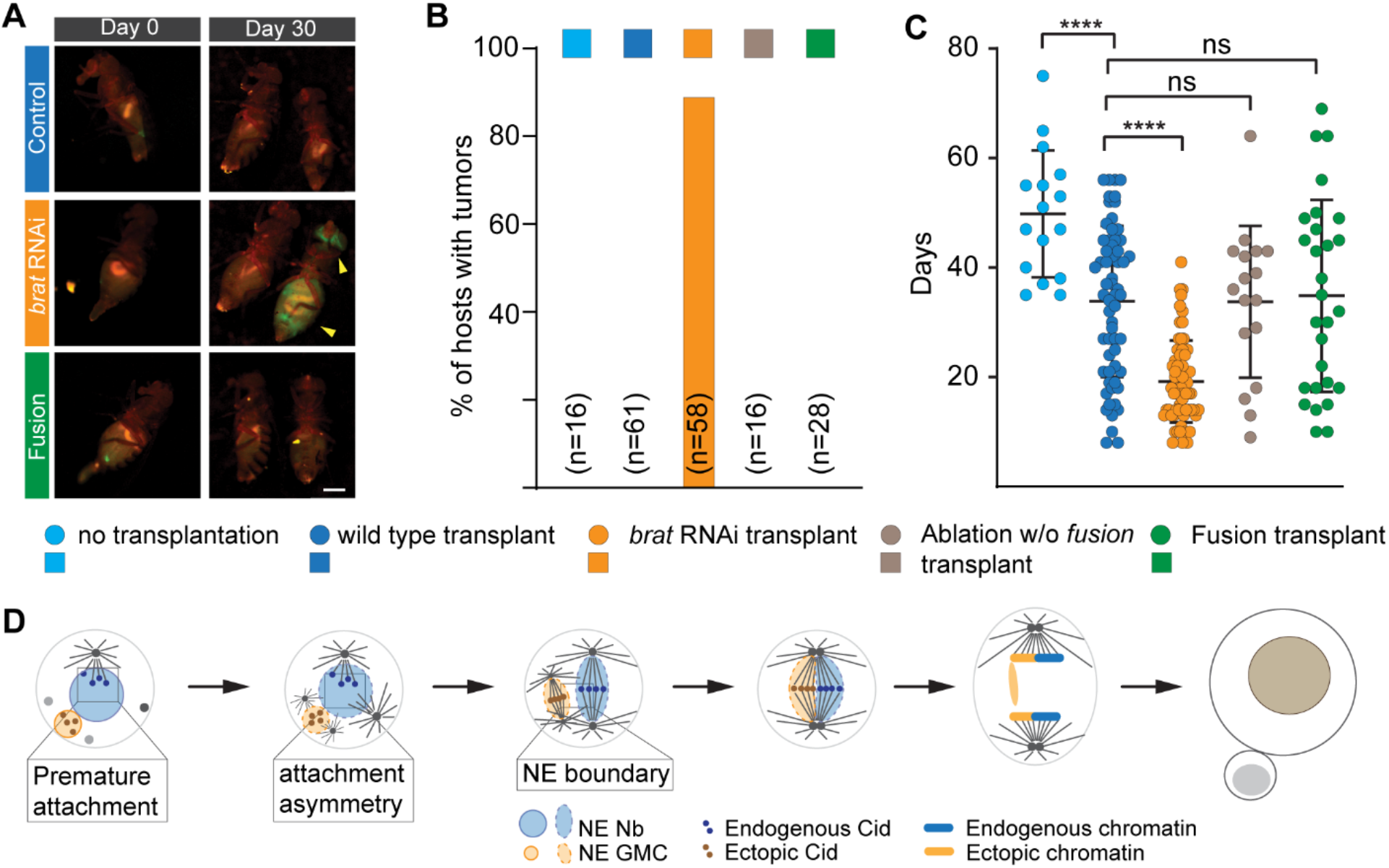
Nb – GMC derived hybrid cells neither form visible tumors nor reduce the lifespan in adult hosts. **(A)** Third instar larval brains expressing His2A::GFP (top row; negative control), co-expressing *brat* RNAi (middle row; positive control) or containing Nb -GMC hybrid cells (bottom row) were transplanted into wild type hosts and imaged at the day of transplantation (day 0) and Day 30. Yellow arrowheads highlight tumors in wild type hosts transplanted with brat RNAi expressing brains. **(B)** Bar graph showing the percentage of host flies containing visible tumors for the indicated grafted samples (colored boxes on top of the graph). **(C)** Scatter plot showing the lifespan of host flies after brain grafts. Grafted tissue either contained no fusions (wild type transplants), expressing *brat* RNAi without fusions (brat RNAi transplants), attempted but unsuccessful fusion (ablation without fusion transplants) and with brains containing hybrid cells (fusion transplants). Host flies without transplanted tissue were plotted for comparison (no transplantation). **(D)** Model: Asymmetric MTOCs capture neuroblast chromatin during interphase, thereby establishing a physical separation between endogenous and ectopic chromatin. Nuclear envelopes form an additional physical barrier. As hybrid cells enter mitosis, the neuroblast centrosomes and introduced ectopic GMC centrosomes nucleate two separate bipolar spindles, aligning their respective chromatin at the metaphase plate. This separation persists through anaphase. Ectopic chromosomes fail to segregate accurately. Error bars correspond to SDs. One way ANOVA. Ns; no signficiance, ****p<0.0001. Scale bar is 1 mm.

## Discussion

Cell – cell fusion can occur under normal physiological conditions and has been implicated in malignancy (Platt and Cascalho, 2019) but how hybrid cells recognize and separate endogenous and ectopic chromosomes during mitosis is not known. Here, we acutely induce cell-cell fusions *in vivo* between Dpn^+^ stem cells and differentiating Pros^+^ GMCs in the developing larval fly brain. We showed that Nb-GMC derived hybrid cells physically separate endogenous neuroblast chromosomes from the introduced ectopic GMC chromosomes and align them independently at the metaphase plate. Chromosome separation is achieved through the formation of two distinct mitotic spindles, which are most likely formed through the canonical centrosome pathway. These dual spindles co-align during metaphase, thereby congressing the two chromosome clusters at the metaphase plate. Endogenous and ectopic chromosomes independently segregate during anaphase, manifested in delayed segregation onset of ectopic chromatin. Although chromosome missegregation is frequent in hybrid cells, potentially leading to aneuploidy, we failed to detect malignant tumor formation when hybrid cell – containing larval brains were grafted into wild type host flies.

We propose that endogenous and ectopic chromosome separation is achieved through an early microtubule-dependent chromosome capture or attachment mechanism that retains endogenous chromosomes in close proximity to the apical neuroblast cortex during interphase. Interphase neuroblasts contain one active MTOC that remains stably anchored close to the apical cell cortex (Rebollo et al., 2007; Rusan and Peifer, 2007). In wild type neuroblasts, chromatin associated Cid is localized in close proximity to the apical MTOC, but Cid’s apical localization is lost upon microtubule-depolymerization or removal of interphase MTOC activity (*cnb* RNAi). Thus, unperturbed wild type neuroblast either retained, or establish microtubule-chromatin connections during interphase. In contrast to yeast, where chromosomes make dynamic attachments to microtubules in G1 (Dorn et al., 2005), this is not the case in other metazoan cells (Maiato et al., 2017). The functional significance of these interphase microtubule-chromatin attachments in neuroblasts are not known but are similar to *Drosophila* male germline stem cells (GSCs), where a single active centrosome connects to chromosomes in prophase, a potential mechanism for biased chromatid segregation (Ranjan et al., 2019).

The geometric separation of endogenous and ectopic chromatin in hybrid cells is further supported by the nuclear envelope, imposing a physical boundary that prevents random chromosome mixing prior to chromosome congression (Figure 6D). In pre-mitotic neuroblasts, chromatin can be connected with centrosomes through the Linc complex (Lee and Burke, 2018), potentially implicating the SUN domain protein Klaroid (Kracklauer et al., 2007) and the KASH-domain protein Klarsicht (Lee and Burke, 2018; Razafsky and Hodzic, 2009) in asymmetric chromatin clustering and the prevention of chromatin mixing during interphase.

The chromatin separation mechanisms described here could be applicable to chromosome separation occurring in the first cleavage after fertilization in insects, arthropods and vertebrates (Kawamura, 2001; Reichmann et al., 2018; Snook et al., 2011). Similarly, biased chromatid and chromosome segregation has been observed in stem cells (Ranjan et al., 2019; Yadlapalli and Yamashita, 2013) and during meiosis, respectively (Akera et al., 2017). Since centromeres have also been found to be confined to specific nuclear locations in many organisms (Muller et al., 2019; Weierich et al., 2003), it will be interesting to see whether microtubule-dependent chromatin attachment provides an alternative mechanism for biased sister chromatid segregation or other important cellular functions.

**Figure 1 – figure supplement 1:**
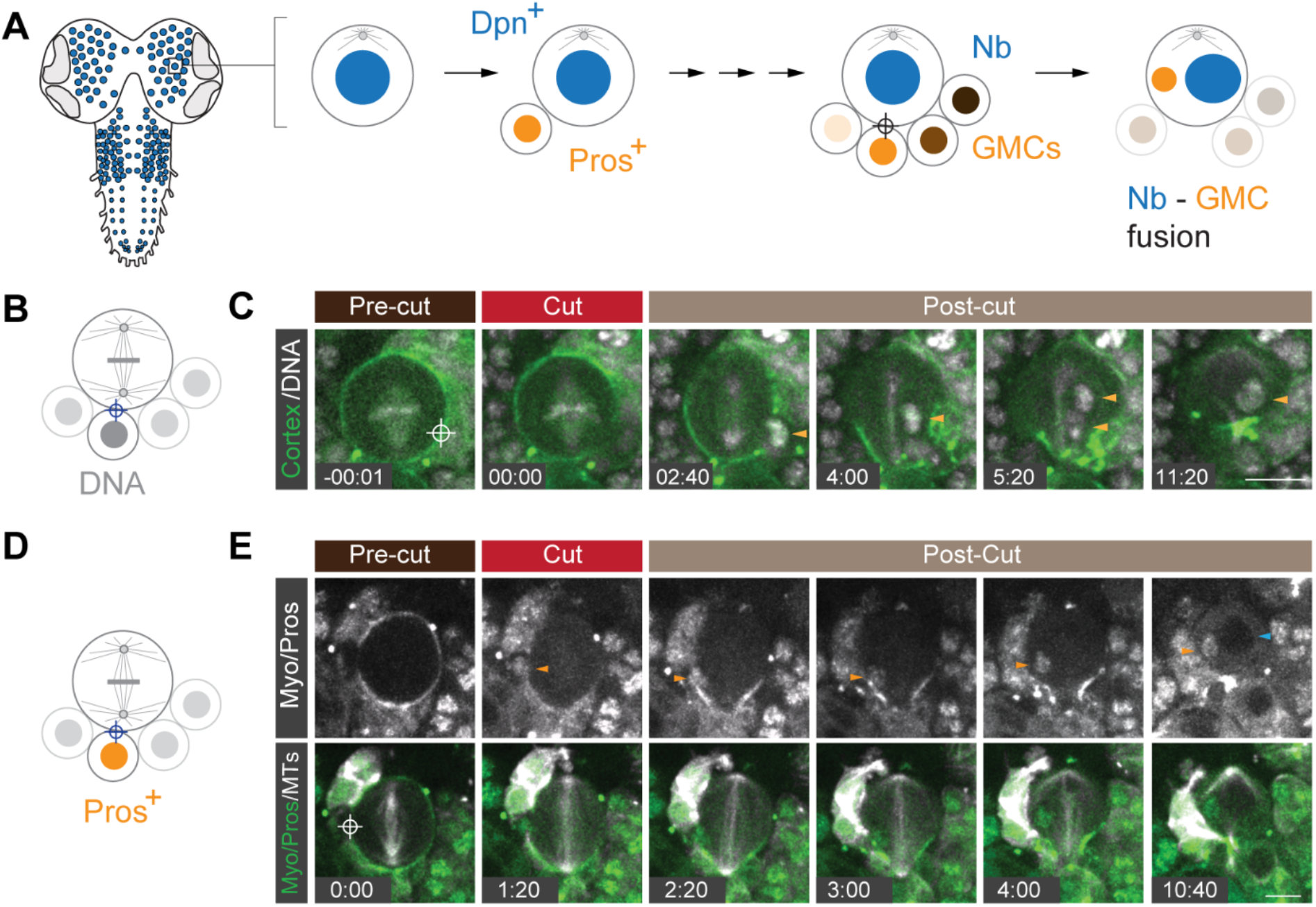
Acute induced Nb – GMC fusion in *Drosophila* larval brains. **(A)** Schematic representation of a third instar *Drosophila* larval brain and neural stem cell division mode. Neural stem cells (neuroblasts (Nbs); blue circles) divide asymmetrically, generating a Prospero-positive, differentiating ganglion mother cell (GMC; Pros^+^) while self-renewing the Dpn^+^ neuroblast. GMCs cluster around Nbs and can be identified based on size and Prospero expression. Acute Nb – GMC fusion can result in hybrid cells containing two molecularly distinct nuclei. **(B)** Experimental outline and **(C)** representative example of a metaphase wild type neuroblast, expressing the cell cortex marker Sqh::EGFP (green), the mitotic spindle marker cherry::Jupiter (white) and the chromatin marker His2A::RFP (white). The site targeted by the ablation laser is highlighted with the white crosshair. The orange arrowhead labels a GMC nucleus moving into the Nb. This hybrid cell successfully completes cytokinesis. **(D)** Schematic and **(E)** representative example of a wild type early anaphase neuroblast expressing Sqh::EGFP (white; top, green; bottom), cherry::Jupiter (white) and Pros::EGFP (white). The orange arrowhead highlights a Pros^+^ GMC nucleus moving into the neuroblast. Cytokinesis completes, creating a hybrid cell containing a Pros^+^ and Pros^-^ nucleus. Time in min:sec; scale bar: 10μm.

**Figure 3 – figure supplement 1:**
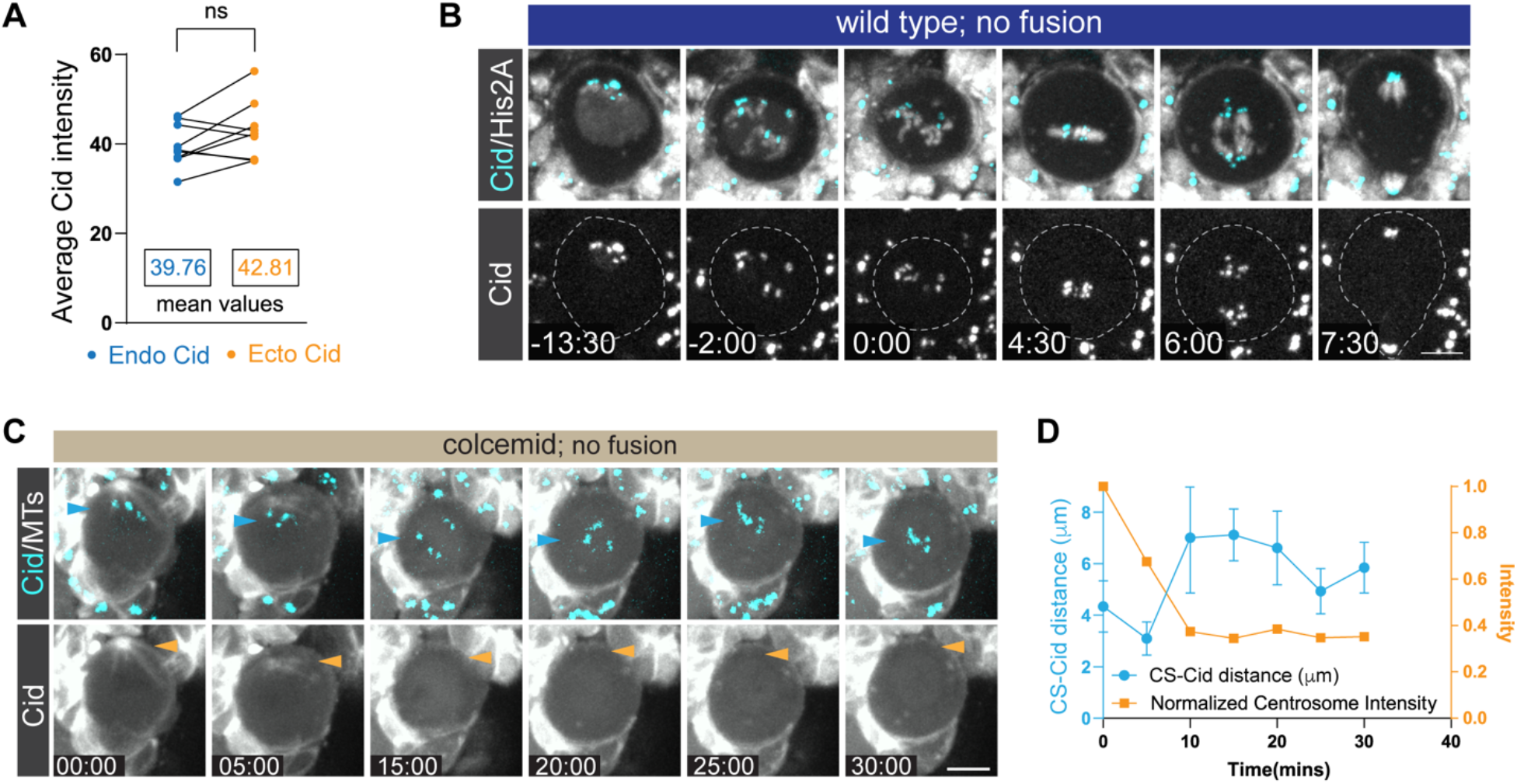
Microtubule-dependent, asymmetric chromatin-centrosome attachments retain chromosomes close to the apical neuroblast cortex during interphase. **(A)** CID intensity measurements of endogenous and ectopic CID in Nb-GMC hybrid cells (p-value, 0.1294). **(B)** Representative wild type neuroblast expressing the canonical histone marker His2A::GFP (white) and Cid::EGFP (cyan; top, white; below). The white dashed line highlights the cell outline. **(C)** Representative example of a wild type neuroblast expressing Cid::EGFP (cyan) and the microtubule marker cherry::Jupiter treated with colcemid. Blue and yellow arrowheads highlight the position of Cid clusters and the disappearing apical MTOC, respectively. **(D)** MTOC intensity and Cid location measurements for the cell shown in (C). Time in mins:secs. Scale bar is 5 μm.

**Figure 4 – figure supplement 1:**
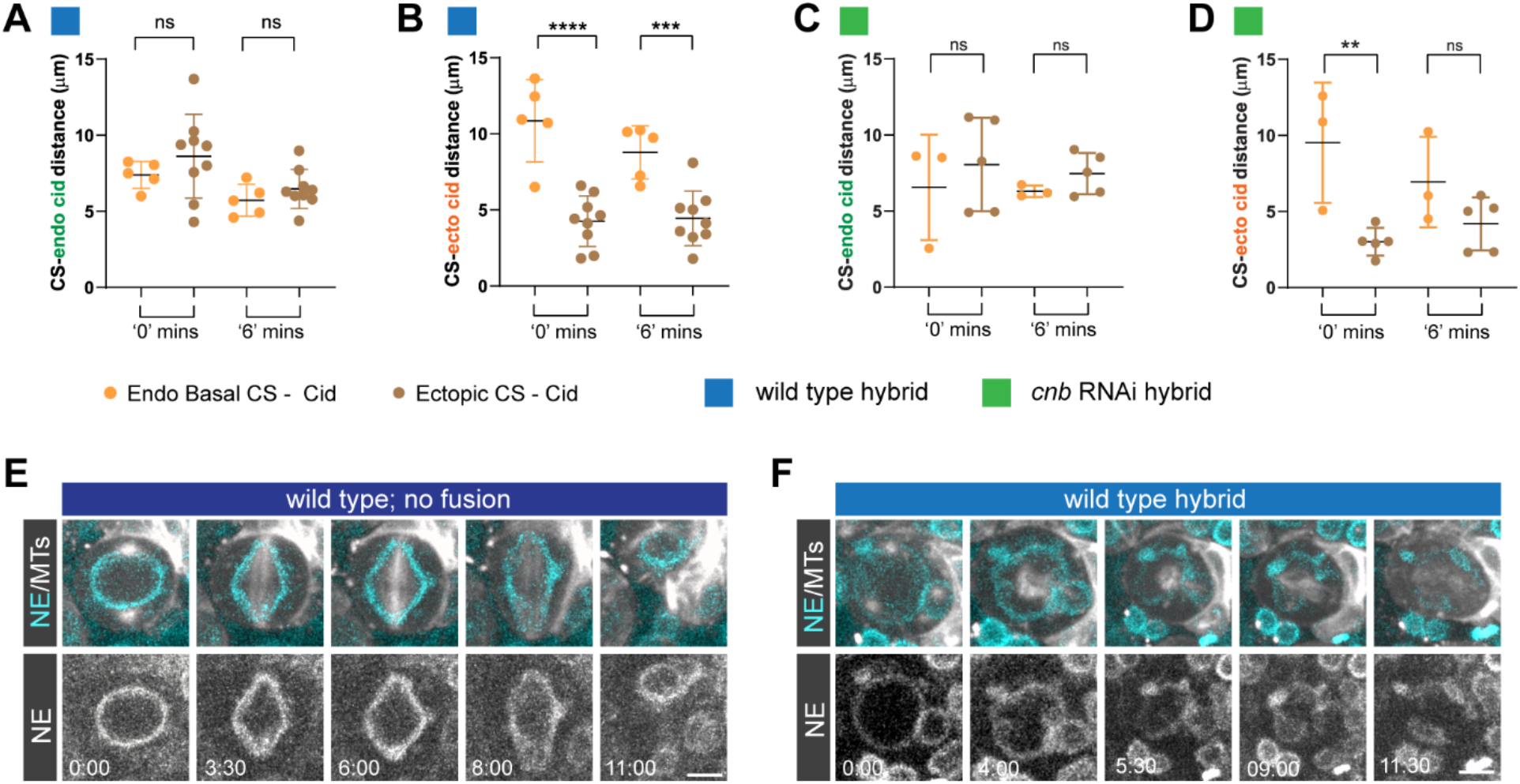
Asymmetric MTOCs and nuclear envelopes separate endogenous and ectopic chromatin in Nb – GMC hybrid cells. **(A)** Averaged distance of basal centrosomes to *endogenous* Cid in wild type or **(C)** *cnb* RNAi expressing neuroblasts. **(B)** Averaged distance of basal centrosomes to *ectopic* Cid in wild type or **(D)** *cnb* RNAi expressing neuroblasts. Measurements are plotted for ‘0’ mins (ectopic centrosome maturation in wild type and *cnb* RNAi expressing hybrids) and 6 mins thereafter (‘6’). Representative image sequence of a wild type **(E)** unfused neuroblasts or **(F)** hybrid cell, expressing the nuclear envelope (NE) marker koi::GFP (cyan; top, white; bottom) and the microtubule marker cherry::Jupiter (white; top). Error bars correspond to SDs. Figure (A-D) twosided unpaired t-test. ns; no significance. ** p<0.01, ***p<0.001, ****p<0.0001. Time in mins:secs. Scale bar is 5 μm.

## Methods

### Fly Strains

Transgenes and fluorescent markers: *worGal4, UAS-mCherry::Jupiter* (Cabernard and Doe, 2009)*; worGal4, UAS-mCherry::Jupiter, Sqh::GFP* (Cabernard et al., 2010)*; His2A::GFP* (Bloomington stock center); *UAS-mCherry::CAAX, UAS-iLID::CAAX:;mCherry (A. Monnard & C. Cabernard; unpublised);* Cid::EGFP (Ranjan et al., 2019); *pUbq-Asl::GFP*(Blachon et al., 2008); *worgal4, UAS-mCherry::Jupiter, Asl::GFP* (this work); *pros::EGFP* (endogenously tagged with CRISPR; this work); *koi::GFP (CB04483)* (Buszczak et al., 2007); *cnb^GD11735^* RNAi line (v28651) (Dietzl et al., 2007).

### Generation of pros::EGFP with CRISPR

Target specific sequences with high efficiency were chosen using the CRISPR Optimal Target Finder (http://tools.flycrispr.molbio.wisc.edu/targetFinder/), the DRSC CRISPR finder (http://www.flyrnai.org/crispr/), and the Efficiency Predictor (http://www.flyrnai.org/evaluateCrispr/) web tools. Sense and antisense primers for these chosen sites were then cloned into pU6-BbsI-ChiRNA (Gratz et al., 2013) between BbsI sites. To generate the replacement donor template, EGFP and 1 kb homology arms flanking the insertion site were cloned into pHD-DsRed-attP (Addgene plasmid #51019) using Infusion technology (Takara/Clontech). Injections were performed in house. Successful events were detected by DsRed-positive screening in the F1 generation. Constitutively active Cre (BDSC#851) was then crossed in to remove the DsRed marker. Positive events were then balanced, genotyped, and sequenced.

### Live cell imaging acute cell-cell fusion

Imaging medium consists of Schneider’s insect medium (Sigma-Aldrich S0146) mixed with 10% BGS (HyClone). Third instar larvae were dissected in imaging medium and the brains were transferred into a μ-slide Angiogenesis or μ-slide 8 well (Ibidi). Live samples were imaged with an Intelligent Imaging Innovations (3i) spinning disc confocal system, consisting of a Yokogawa CSU-W1 spinning disc unit and two Prime 95B Scientific CMOS cameras. A 60x/1.4NA oil immersion objective mounted on a Nikon Eclipse Ti microscope was used for imaging. Live imaging voxels are 0.22 × 0.22 × 0.75-1 □m (60x/1.4NA spinning disc).

Neuroblast-GMC fusions were induced using a 3i Ablate! ablation system, consisting of a 532nm pulsed laser. We used a pulse width of 7 ns, targeting the membrane interface between the neuroblast and the adjacent GMC.

### Colcemid treatment

Dissected brains were incubated with Colcemid (Sigma) in live imaging medium at a final concentration of of 25 μgmL·^1^.

### Transplantation experiments

Brain lobes containing the hybrid cells expressing *His2A::GFP* and *worGal4, UAS-mCherry::Jupiter* were transplanted into 3 to 4 day old, well fed adult *w^1118^* female host flies as described previously (Rossi and Gonzalez, 2015). Custom made needles were prepared from Narishige GD-1 glass capillaries using a Narishige, needle puller. Injection needles were shaped with forceps to have a smooth, 45° opening. Transplanted flies were transferred into fresh vials each day for the first three days, followed by biweekly flipping. The tumor growth was monitored and recorded with a Leica MZ FLIII fluorescence stereomicroscope.

### Image processing and measurements

Live cell images were processed using imaris x64 8.3.1 and image J.

For angle and distance measurements, the coordinates for the two spindle poles were determined in Imaris. From these coordinates, angles and distances between spindles were derived based on the calculations outlined below.

*Angle between spindles*: 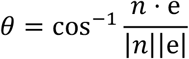

Dot product: 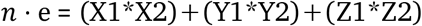

Magnitude of vectors: 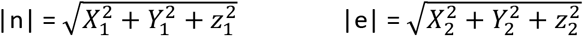

Where **n** corresponds to the spindle vector: **n** (x1, y1, z1) = (N1-N1 ‘, N2-N2’, N3-N3’) and **e** to the ectopic spindle vector: **e** (x2,y2,z2) = (E1-E1’, E2-E2’, E3-E3’)

N1, N2, N3 and N1’, N2’ and N3’ are coordinates of the two poles of the Nb spindle. Similarly, E1, E2, and E3 and E1’, E2’ and E3’ are coordinates of the ectopic spindle poles.

### Distance between spindle vectors

The midpoints of the two spindle vectors are calculated from coordinates of the poles on either side of the respective spindle. This is followed by calculating the distance between these midpoints.

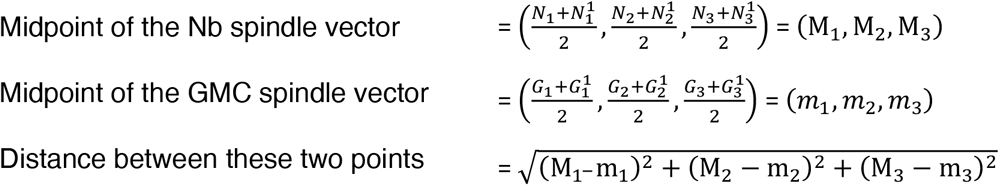

### Centrosome – Cid distance

The centrosome (CS) – Cid distance was calculated using Cid and CS coordinates.

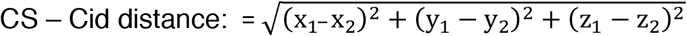

Where x_1_,y_1_,z_1_ correspond to CS and x_2_,y_2_,z_2_ to Cid coordinates, respectively.

Plotted values correspond to averaged values of all CS – Cid punctae distances and the corresponding standard deviations.

0 and 6 mins corresponds to the appearance of the basal centrosome and 6 minutes thereafter. ‘0’ and ‘6’ mins corresponds to the appearance of the ectopic centrosome and 6 minutes thereafter.

### Statistical analysis

Statistical analysis was performed using Graphpad prism 8. Statistical significance was determined using paired or unpaired t-test and one-way ANOVA. Significance was indicated as following: *; p<0.05, **;p<0.01, ***;p<0.001,****p<0.0001, ns; not significant. Exact p values and complete statistical information can be found in Extended data table 1.

Measurements were taken from distinct samples and from several independent experiments.

## Acknowledgements

We thank Xin Chen for fly stocks, David Salvador Garcia for generating the Pros::EGFP transgenic line, Sue Biggins and members of the Cabernard laboratory for helpful discussions and comments. This work was supported by the National Institutes of Health (1R01GM126029-03) and a Research Scholar grant from the American Cancer Society (130285-RSG-16253-01-CSM). Stocks obtained from the Bloomington Drosophila Stock Center (NIH P40OD018537) and from the Vienna Drosophila Resource Center (VDRC).

## Author contributions

This study was conceived by B.S., N.L., and C.C. C.R provided some conceptual ideas early on. Technical feasibility and was demonstrated by C.R & C.C.

B.S and N.L performed all the experiments with significant help from C.S.

B.S, N.L., C.S., and C.C analyzed the data.

B.S and C.C. wrote the manuscript.

## Data availability statement

The authors declare that all data supporting the findings of this study are available within the paper and its supplementary files. Source data are available upon request.

## Supplementary Table 1: Statistical information

Complete statistical information for the data shown in the corresponding figures.

## Video legends

**Video 1: Wild type neuroblast division; related to Figure 1b**

Wild type control (unfused) neuroblast expressing the microtubule binding protein Cherry::Jupiter (white) and the canonical Histone marker His2A::GFP (cyan). Time scale is h:mm:ss and the scale bar is 5 μm.

**Video 2: Wild type hybrid cell; related to Figure 1b**

Wild type hybrid cell derived from a neuroblast – GMC fusion *in vivo*, expressing the canonical Histone marker His2A::GFP (white in single channel; cyan in merge) and the microtubule binding protein Cherry::Jupiter (white). The blue and orange arrows mark endogenous and ectopic chromatin, respectively. Time scale is h:mm:ss and the scale bar is 3 μm.

**Video 3: Wild type hybrid cell; related to Figure 2b**

Wild type hybrid cell derived from a neuroblast – GMC fusion *in vivo*, expressing the canonical Histone marker His2A::GFP (white in single channel; cyan in merge) and the microtubule binding protein Cherry::Jupiter (white). The blue and orange arrows mark endogenous and ectopic chromatin, respectively. Time scale is h:mm:ss and the scale bar is 2 μm.

**Video 4: Wild type neuroblast division; related to Extended Data Figure 2b**

Wild type control (unfused) neuroblast, expressing the microtubule binding protein Cherry::Jupiter (white) and the centromere-specific H3 variant Cid::EGFP (Cyan). Purple and yellow arrows point to the apical and basal centrosome, respectively. The blue arrow refers to moving Cid clusters. Time scale is h:mm:ss and the scale bar is 1 μm.

**Video 5: Wild type neuroblast division; related to Extended Data Figure 2f**

Wild type control (unfused) neuroblast, expressing the membrane marker mCherry::CAAX (white), the canonical Histone marker His2A::GFP (white) and Cid::EGFP (Cyan). The green arrow points to Cid clusters. Time scale is h:mm:ss and the scale bar is 5 μm.

**Video 6: Wild type neuroblast exposed to Colcemid; related to Extended Data Figure 2g**

Wild type control (unfused) neuroblast exposed to the microtubule depolymerizing drug Colcemid, expressing the membrane marker mCherry::CAAX (white), the microtubule binding protein Cherry::Jupiter (white) and Cid::EGFP (Cyan). The yellow arrow points to the apical centrosome, the blue arrow to Cid clusters. Time scale is h:mm:ss and the scale bar is 1 μm.

**Video 7: Cnb RNAi expressing neuroblast; related to Extended Data Figure 2i**

*cnb* RNAi expressing (unfused) neuroblast, co-expressing the microtubule binding protein Cherry::Jupiter (white) and Cid::EGFP (Cyan). The orange and blue arrow points to the apical and basal centrosome, respectively. The green arrow highlights Cid clusters. Time scale is h:mm:ss and the scale bar is 1 μm.

**Video 8: wild type hybrid cell; related to Extended Data Figure 3a**

Wild type hybrid cells expressing the microtubule binding protein Cherry::Jupiter (white) and Cid::EGFP (white in single channel; Cyan in merge). The blue and orange arrow highlights endogenous and ectopic Cid clusters, respectively. Time scale is h:mm:ss and the scale bar is 1 μm.

**Video 9: Cnb RNAi expressing hybrid cell; related to Extended Data Figure 3d**

*cnb* RNAi expressing hybrid cell, expressing the microtubule binding protein Cherry::Jupiter (white) and Cid::EGFP (white in single channel; Cyan in merge). The blue and orange arrow highlights endogenous and ectopic Cid clusters, respectively. The yellow arrow indicates mixing of endogenous and ectopic Cid clusters. Time scale is h:mm:ss and the scale bar is 1 μm.

**Video 10: wild type hybrid cell; related to Figure 4e**

Wild type hybrid cell, expressing the canonical Histone marker His2A::GFP (white). The blue and orange arrow highlights endogenous and ectopic chromosomes, respectively. The magenta arrowhead highlights the fate of missegregated chromosomes. This hybrid cell forms a heterokaryon. Time scale is h:mm:ss and the scale bar is 1 μm.

